# Increasing cell-free gene expression yields from linear templates in *Escherichia coli* and *Vibrio natriegens* extracts by using DNA-binding proteins

**DOI:** 10.1101/2020.07.22.214221

**Authors:** Bo Zhu, Rui Gan, Maria D. Cabezas, Takaaki Kojima, Robert Nicol, Michael C. Jewett, Hideo Nakano

## Abstract

In crude extract-based cell-free protein synthesis (CFPS), DNA templates are transcribed and translated into functional proteins. Although linear expression templates (LETs) are less laborious and expensive to generate, plasmid templates are often desired over PCR-generated LETs due to increased stability and protection against exonucleases present in the extract of the reaction. Here we demonstrate that addition of a dsDNA-binding protein to the CFPS reaction, termed single-chain Cro protein (scCro), achieves terminal protection of LETs. This CroP-LET (sc<u>Cro</u>-based <u>P</u>rotection of <u>LET</u>) method effectively increases sfGFP expression levels from LETs in *Escherichia coli* CFPS reactions by 6-fold. Our yields are comparable to other strategies that provide chemical and enzymatic DNA stabilization in *E. coli* CFPS. Notably, we also report that the CroP-LET method successfully enhanced yields in CFPS platforms derived from non-model organisms. Our results show that CroP-LET increased sfGFP yields by 18-fold in the *Vibrio natriegens* CFPS platform. With the fast-expanding applications of CFPS platforms, this method provides a practical and generalizable solution to protect linear expression DNA templates.

## Introduction

Crude extract-based cell-free protein synthesis (CFPS) provides a practical *in vitro* approach for protein expression. By combining the translation machinery present in the cell extract with additional enzymes, co-factors, amino acids, tRNAs, energy sources, and other small molecules in a test tube (Carlson et al., 2012), this approach can produce functional proteins within hours from a plasmid or linear expression template (LET). While CFPS conditions may vary according to the unique transcription/translation requirements for the synthesis of individual proteins, these reactions can be tailored by optimizing concentrations of the CFPS reagents. As such, CFPS has enabled a wide range of applications in directed evolution, synthetic biology, glycoscience, metabolic engineering, and education (Des Soye et al., 2019; Harris and Jewett, 2012; Karim and Jewett, 2016; Kightlinger et al., 2019; Lin et al., 2020; Martin et al., 2018; Silverman et al., 2020; Stark et al., 2019; Thavarajah et al., 2020). LETs offer a unique advantage over their circular counterparts because extensive plasmid construction steps are not required. However, LET degradation by exonucleases present in the cell extract still remains a significant challenge that prevents the wide use of LETs for CFPS (Michel-Reydellet et al., 2005).

Frequently, exonucleases present in crude cell extracts are responsible for LET degradation in the CFPS reaction (Hoffmann et al., 2002; Michel-Reydellet et al., 2005). Previous works in *Escherichia coli-based CFPS* have addressed this problem by chemically modifying LETs (such as introducing unnatural 3’-end or 5’-end adenosine and addition of loop ends) (Hoffmann et al., 2002), addition of GamS to inhibit RecBCD (Sun et al., 2014), and supplementing chi-site DNA oligos to the CFPS reaction(Marshall et al., 2017). However, these strategies may not be applicable to CFPS systems derived from non-model organisms. For example, the LET-stabilization effect was not significantly observed in *V. natriegens*-based CFPS in the presence of GamS, where a yield of superfolder green fluorescent protein (sfGFP) at 0.02 mg/mL from 30 nM LET was detected (Wiegand et al., 2018), Another common strategy involves engineering strains to remove proteins and enzymes that degrade linear templates. Previously, the Swartz group developed a genetically modified *E. coli* strain (A19) to stabilize the DNA template by removing endA (endonuclease I) and RecBCD from the genome. While these strategies showed increased stability of the DNA template and improved protein production yields for *E. coli* CFPS (Michel-Reydellet et al., 2005), these methods require strain engineering efforts that might not port to all organisms and are not off-the-shelf solutions (Kelwick et al., 2016; Li et al., 2017; Martin et al., 2017; Wang et al., 2018).

To address this issue, we identified a DNA binding protein that provides increased stability to LETs against exonuclease degradation. This dsDNA-binding protein, termed single-chain lambda Cro repressor (scCro), directly protects the free termini of the LET with a dsDNA-binding protein by sterically blocking the progressive degradation caused by various exonucleases in the crude cell extract (**Figure 1A**). In this CroP-LET (sc<u>Cro</u>-based <u>P</u>rotection of <u>LET</u>) method, the scCro specifically binds to a 17-bp dsDNA operator recognition consensus sequence (ORC) at high affinity with a *K*_D_ in the range of 4 pM to 1.8 nM (Jana et al., 1998; Kojima et al., 2018; Nilsson and Widersten, 2004). The three-dimensional structure revealed that binding specificity is accomplished by direct hydrogen-bonding and van der Waals interactions between the protein and the exposed bases of both strands (**Figure 1B**) within the major groove of the DNA (Albright and Matthews, 1998; Nilsson and Widersten, 2004). Moreover, the binding affinity ratio between the 17-bp ORC and non-specific DNA with scCro is 10,000:1, which indicates a low probability of interference with any other sequence than the ORC (Kim et al., 1987). Leveraging the high affinity and specificity, scCro has been successfully utilized for protein immobilization in microbead display (Kojima et al., 2016; Zhu et al., 2015) and to control the spatial arrangement of enzymes on DNA scaffolds for designing reaction cascades (Kojima et al., 2018).

**Figure 1.**
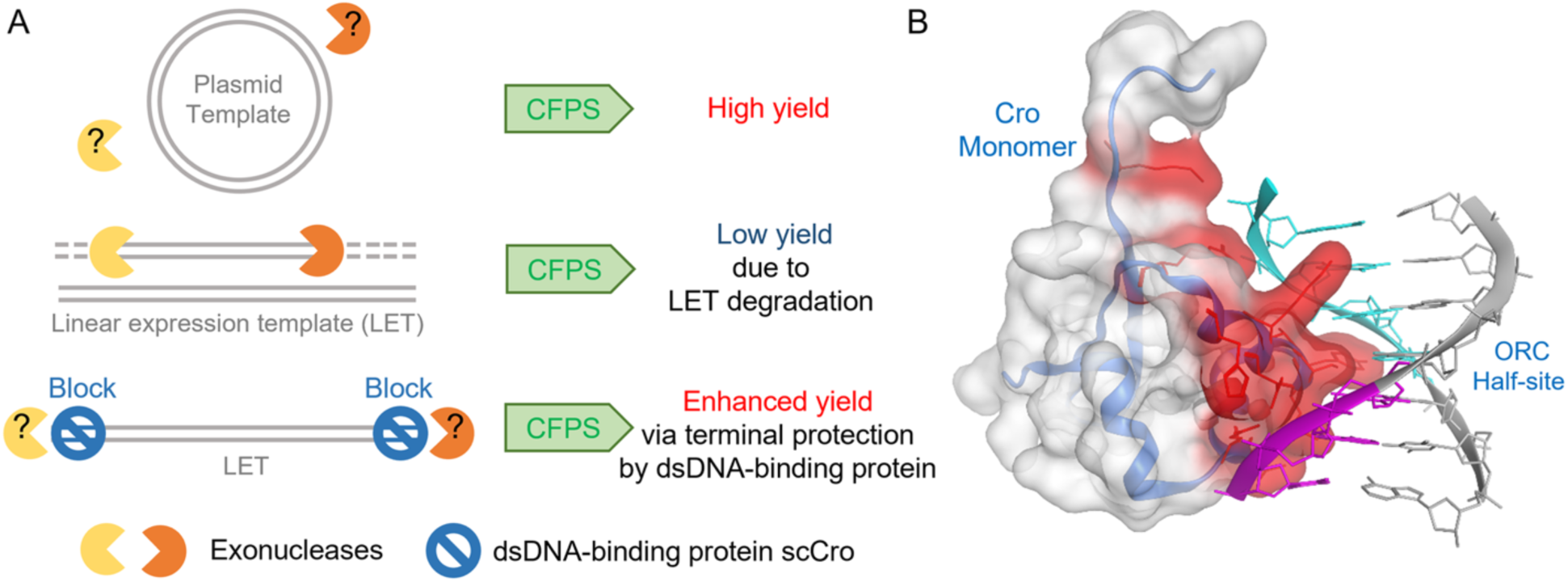
LET terminal protection prevents exonuclease degradation and increases protein yields in CFPS. **(A)** Proposed mechanism of LET terminal blocking by scCro. **(B)** Three-dimensional structure of lambda Cro repressor monomer and an operator recognition consensus sequence (ORC) half-binding site complex (PDB code: 6CRO). The surface of the Cro monomer is shown in transparent mode. The blue ribbon represents the main chain of the Cro protein. The residues interacting with the ORC half-site are colored in red. The grey ribbon indicates the dsDNA sequence of the ORC half-site TATCACC. The nucleotides interacting with the Cro protein were labeled in magenta or cyan for the sense and antisense strand, respectively.

In this study, we develop a generalizable, easy-to-use method called CroP-LET for improving the stability of LETs using scCro. We demonstrate that this approach can be applied to both *E. coli* and *Vibrio natriegens* CFPS platforms. First, we show that sfGFP yield in *E. coli* CFPS increased by 6-fold when the LET was terminally protected by scCro thus achieving yields comparable to RecBCD inhibition with GamS. Second, we show host-versatility of this approach by applying the scCro-based protection method to non-model CFPS platforms, specifically for *V. natriegens*. Significantly, the yield sfGFP expression from LET in the *V. natriegens* CFPS system increased by up to 18-fold to 0.3 mg/mL. Given the accelerated discovery of novel CFPS systems, the CroP-LET method presented here provides a rapid and cross-species compatible solution for protecting LETs. We anticipate that our results will accelerate the characterization of unknown gene functions (Salzberg, 2019) and enable the rapid synthesis of template libraries (Shrestha et al., 2014) for *in vitro* directed evolution studies (Ueno et al., 2012; Zhu et al., 2015), among other applications by utilizing low-cost commercially synthesized LETs in high-throughput CFPS-based screening platforms.

## Results and Discussion

### Terminal protection of LET increases linear DNA stability in *E. coli* CFPS

We first generated two LETs to probe whether scCro could be used for effective terminal blocking in CFPS (**Figure 2A**). LET1 was designed as a linear counterpart to the plasmid template, with buffer sequences added to each end. LET2 comprises the same elements of LET1, but also includes the addition of terminal ORC binding sites that could be used to assess the effect of scCro-based terminal blocking. To confirm LET protection by scCro, we initially tested the susceptibility of LET2 degradation in presence of *E. coli* exonuclease RecBCD (**Figure 2B**). As expected, RecBCD degraded the unprotected LET2 rapidly, within 15 min of incubation. We also observed that terminal blocking with scCro protected 65 % of the LET2 from degradation after 15 min of incubation with RecBCD, while only 3 % of the LET2 remained full length in the absence of scCro (**Figure 2C**). These data suggest that scCro sterically hinders RecBCD association with the LET2 ORC and prevents degradation.

**Figure 2.**
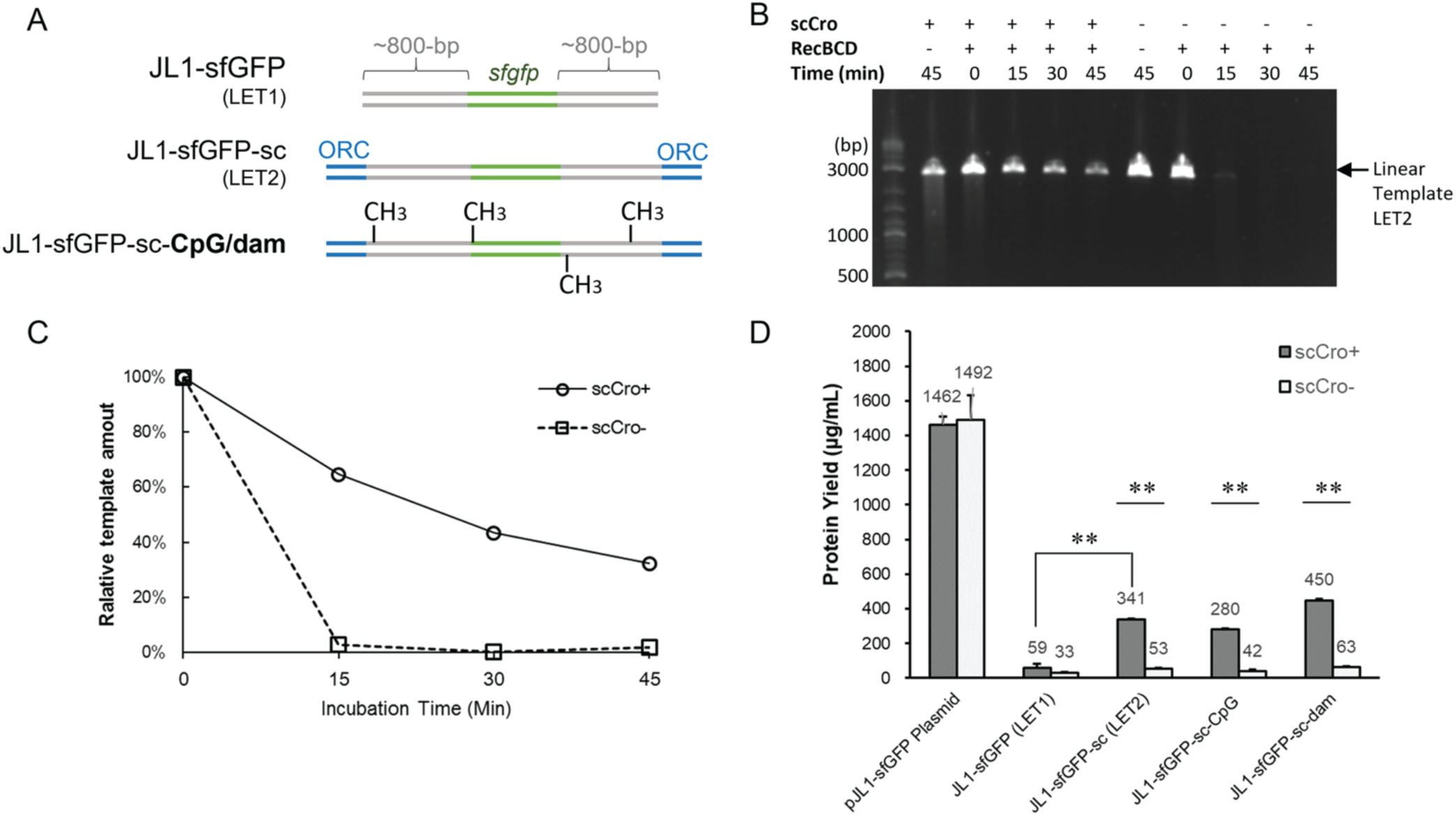
Effect of scCro-based protection of LETs on sfGFP yields in *E. coli* CFPS. **(A)** LET sequences used in this study. *sfgfp* corresponds to the sfGFP operon. ORC refers to the scCro-binding site. JL1-sfGFP (LET1): the linearized full-length plasmid pJL1-sfGFP with buffer regions at each end marked in gray generated by PCR; JL1-sfGFP-sc (LET2): the JL1-sfGFP sequence with buffer regions marked in gray and one ORC at each end of the LET marked in blue; JL1-sfGFP-sc-CpG/dam: the JL1-sfGFP-sc (LET2) template with further methylation by CpG or *dam* methyltransferases. **(B)** Degradation of linear template JL1-sfGFP-sc by RecBCD in the presence of scCro. **(C)** Gel quantification analysis of the scCro-protected LET using ImageJ. **(D)** Yields of sfGFP using scCro-protected and methylated LETs in CFPS. scCro+ indicates the addition of scCro to DNA templates before the CFPS reactions while scCro-indicates that scCro was absent. The mean and standard deviations are shown for N=4. P-values were determined using the Welch’s t-test; **P < 0.0005.

With LET2 having demonstrated stabilization in the presence of exonucleases, we next tested the impact of scCro protected DNA in CFPS. sfGFP expression levels were compared in the presence and absence of scCro in the CFPS reaction (**Figure 2D**). Cell extract was prepared from *E. coli* strain *C321*.*ΔA*.*759*, in which DNA endonuclease I *(endA−*) and ribonuclease E (*rne−*)(Martin et al., 2018) were inactivated to minimize the effect endonuclease and RNA degradation on the final yield of sfGFP. As a control, addition of scCro to a plasmid-based CFPS reaction did not have a significant effect on sfGFP yields (**Figure 2D**) and the observed yields are comparable to our previous published results.^23^ When the LET does not contain any scCro-binding sites—as is the case in the LET1 sequence—the LET is still susceptible to degradation resulting in poor sfGFP yields ∼ 0.05 mg/mL. In contrast, addition of scCro to the CFPS reaction expressing the LET2—which contains terminal binding sites for scCro—resulted in a 6-fold protein yield improvement over the condition without scCro (**Figure 2D**).

We also compared our approach with DNA methylation strategies used to stabilize DNA templates. We observed methylation of LET2 using *dam* methyltransferase further increased sfGFP yield by 32 %, while modification with CpG methyltransferase reduced sfGFP yield by 18 % in absence of scCro (**Figure 2D**). Furthermore, the synergistic effect of LET protection and DNA methylation did not enhance sfGFP expression significantly. From these results, we conclude that the effect of LET methylation on sfGFP production is not significant. Therefore, unmethylated LETs were used in the experiments described below.

Another common approach to prevent LET degradation in *E. coli* CFPS is to supplement the reaction with exonuclease V inhibitor protein GamS. Our results show that addition of either GamS or scCro resulted in comparable levels of sfGFP expression (**Figure S1**). While GamS is known to fully inhibit RecBCD exonuclease activity, we hypothesize that scCro offers a broader range of protection against degradation by exonucleases—including RecBCD—present in the extract. For this reason, we tested the synergistic effects of adding both GamS and scCro, however, we found no significant synergistic effect (**Figure S1**). Next, we monitored the blocking effect of scCro at a range of concentrations 5 – 40 μg/mL (corresponding to 0.28 – 2.2 μM) in presence of 8 nM LET and found that degradation can be inhibited at scCro concentrations as low as 0.28 μM (**Figure S2**). Previously, a concentration of 1 μM of GamS with 2 nM LET was sufficient to prevent degradation and improve expression of deGFP in *E. coli* CFPS (Sun et al., 2014). In this work, our results indicate that sufficient protection of LET can be achieved at relatively lower scCro concentrations compared to GamS because of the high affinity between scCro and the ORC (Jana et al., 1998; Kojima et al., 2018; Nilsson and Widersten, 2004).

### Synergetic effects between scCro-based protection and buffer sequences

We next evaluated if the DNA bases upstream and downstream of the linear *sfgfp* operon are required for CroP-LET mediated protection. Previously, DNA sequences flanking the *sfgfp* operon (*i*.*e*., buffer sequences), which originate from the amplification template used to make the LET, have been shown to sufficiently alleviate some LET degradation in *E. coli* CFPS (Sun et al., 2014). To test whether introducing buffer sequences (∼ 800-bp buffer sequence at both ends) between the *sfgfp* operon and ORC binding sites would provide additional stability of LET2 alone, we compared LET2 to LET3, a counterpart linear template to LET2 which has no buffer sequences, and tested these LETs in CFPS with and without scCro (**Figure 3A**). The presence of buffer sequence, as show in LET2, contributed to ∼7-fold yield increase in CFPS regardless of the presence of scCro protein (**Figure 3B**); this demonstrates that a long buffer sequence is able to protect LETs. When scCro was present, we observed a ∼6-fold improvement in sfGFP yields for both LET2 and LET3 (**Figure 3B**), which implies that the effect of scCro protection is independent of the buffer sequence. These results suggest there is a synergetic effect between the scCro-based protection and the presence of the buffer sequences. This observation could be because buffer sequences allow scCro to bind the 5’-end of the LETs without inhibiting transcription. Previously, it has been shown that phosphorothioate bond modifications can inhibit exonucleases (Putney et al., 1981; Yang et al., 2007) and introducing this type of modification potentially stabilizes LETs. We therefore tested whether adding phosphorothioate modifications (LET5) affected protein synthesis from LET. However, same as the LET4, LET5 only produced a low amount (5-10 µg/mL) of sfGFP.

**Figure 3.**
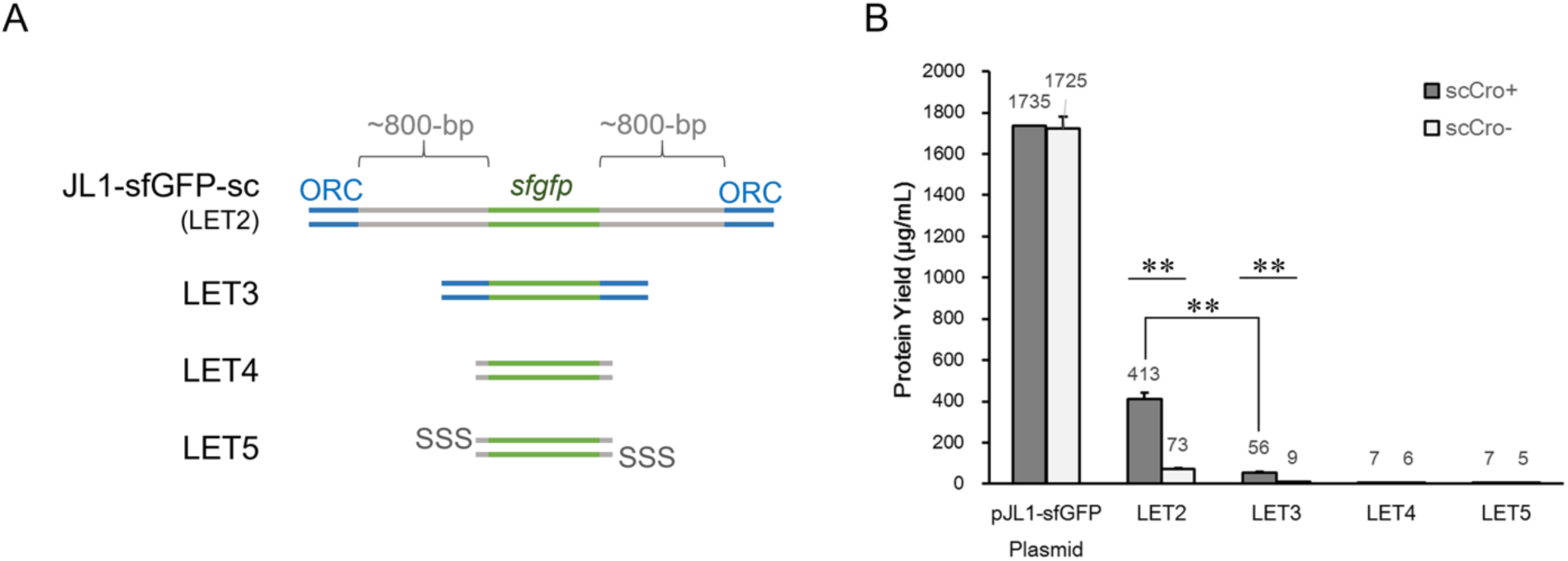
Synergetic effects between scCro-based protection and buffering sequences. **(A)** Diagram describing LETs used in this study. The regions colored in green correspond to the operon of *sfgfp*. The regions in blue ORC refer to the scCro-binding site. JL1-sfGFP-sc (LET2): the linearized full-length plasmid pJL1-sfGFP generated by PCR with one ORC at each end, and the PCR-amplified gray regions flanking outside the sfGFP operon are referred as the buffer sequences in this study. LET3: short LET without buffer sequences, only ORC regions in blue; LET4: short LET without buffer sequence and ORC; LET5: short LET (contains neither buffer sequence nor ORC) with three 5’ phosphorothioate bonds (SSS) at each end. **(B)** sfGFP yield from different LETs expressed in CFPS. scCro+ indicates that scCro was added to samples before the CFPS reactions. scCro-indicates that scCro was not added to samples. The mean and standard deviations are shown (N=4). P-values were determined using the Welch’s t-test; **P < 0.0005.

### scCro prevents LET degradation in the *V. natriegens* CFPS system

Recently, a novel high-yielding CFPS platform derived from *V. natriegens* was developed (Des Soye et al., 2018; Wiegand et al., 2018). This non-model organism has a fast doubling time (∼10 min) and the potential to serve as a robust alternative for the cell-free production of proteins, commodity chemicals and other synthetic biology applications (Des Soye et al., 2018; Failmezger et al., 2018; Wiegand et al., 2018). While sfGFP yields up to ∼1.6 mg/mL have been achieved in the *V. natriegens* CFPS platform from a plasmid template, only yields ∼0.02 mg/mL of sfGFP from a 30 nM LET have been reported, thus a solution is required in order to improve protein yields from LET-based systems (Wiegand et al., 2018). Given that scCro prevents degradation of LETs in *E. coli* CFPS, we decided to explore scCro as an alternative solution to improve yields from LETs in the *V. natriegens* CFPS system (**Figure 4**). Similar to our *E. coli* experiments, we incubated scCro with LETs prior to the CFPS reaction. In both *E. coli* and *V. natriegens* CFPS platforms, we observed sfGFP expression levels comparable to previous reports and addition of scCro did not have a significant to impact sfGFP expression. Without ORC binding sites, expression from LET1 yielded less than 10% of sfGFP in the *V. natriegens* system compared to the *E. coli* CFPS platform; this result points out that nucleases in the *V. natriegens* CFPS are more active. Interestingly, we observed an 8-fold increase corresponding to 0.03 mg/mL of overall sfGFP production from 8 nM LET2 (**Figure 4**). Although this amount is 10% lower (∼0.4 mg/mL) than the yield observed in *E. coli* CFPS, we hypothesize that endogenous nucleases specific to *V. natriegens* are responsible for LET degradation. We observed that sfGFP yield increased proportionally with the concentration of LET2. In this way, we achieved sfGFP yields of up to 0.324 mg/mL with 64 nM of LET2, while keeping the concentration of scCro constant at 2 μM for all conditions (**Figure 4**). We found that scCro-based protection became more significant at higher LET concentrations. With 64 nM of LET2 in the CFPS reaction, scCro-protected template showed an 18-fold improvement in sfGFP yield over the non-protected template, thus 2 μM of scCro is sufficient to protect up to 64 nM of the LET. While further increasing the DNA concentration is expected to enhance sfGFP yield, we anticipate that it would be challenging to assemble CFPS reactions at concentrations beyond 64 nM of LET template due to a volume limitations. That said, protein yields of ∼0.3 mg/mL are sufficient to enable a wide range high-throughput screening applications, including directed evolution of enzymes, prototyping of synthetic biological circuits, and optimization of metabolic engineering pathways.

**Figure 4.**
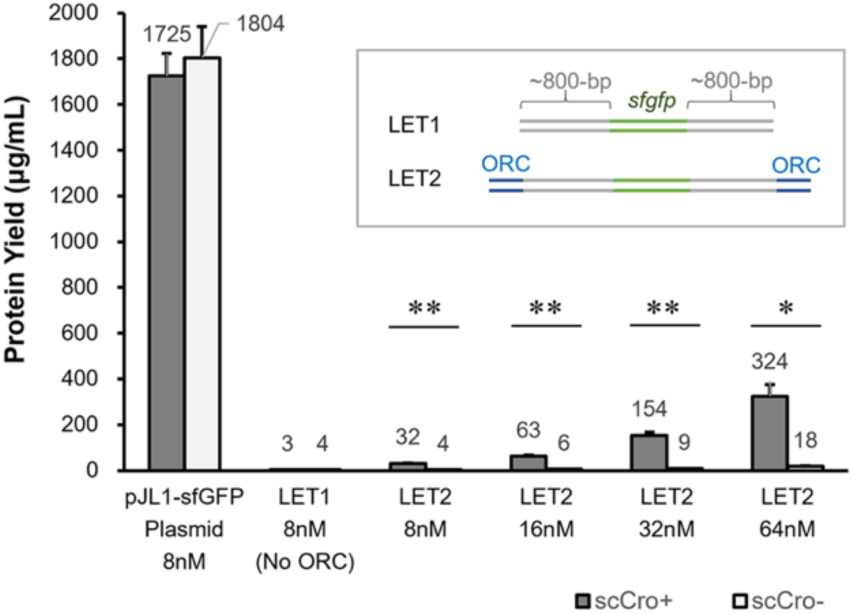
scCro-based protection of LETs boosts sfGFP yields in the *V. natriegens* CFPS system. LET1 and LET2 are the same templates described in Figure 2A. scCro+ indicates that scCro was added to the samples before the CFPS reactions. scCro-indicates that scCro was not added to templates. The mean and standard deviations are shown (N=4). P-values were determined using the Welch’s t-test; *P < 0.005, **P < 0.0005.

### Conclusions

In this study, we developed a novel method termed CroP-LET that increases protein yields in CFPS systems by terminally blocking degradation of linear expression templates with the dsDNA-binding protein scCro. This method only requires addition of a 17-bp scCro-binding site sequence at the 5’-end of the primers used to amplify LETs. The scCro protein can be easily obtained via overexpression in *E. coli* followed by a one-step His-tag purification. Equally important, this method is successful both in the *E. coli* and *V. natriegens* CFPS systems, providing a generalizable solution that is independent of the hosts’ genetic and biochemical background and can be applied CFPS systems derived from non-model organisms. Taken together, our method provides a simple and cross-species tool that will enable non-model CPFS applications.

## Materials and Methods

### Reagents and buffers

Chemicals were purchased from Sigma-Aldrich unless designated otherwise. DNA polymerase (Phusion), CpG methyltransferase (M.SssI), dam methyltransferase, RecBCD (exonuclease V), were purchased from New England Biolabs (NEB). T7 RNA polymerase was prepared as previously described.(Swartz et al., 2004) Plasmids were extracted using a Plasmid Miniprep Kit (Omega Bio-Tek). All DNA oligonucleotides were purchased from Integrated DNA Technologies, Inc (IDT) (**Table S1**).

### Linear expression templates (LETs) preparation

To test the effect of LETs in CFPS, various LET sequences were prepared by PCR amplification (**Figure 2A and 3A**). LET1 was obtained by amplifying the full length of the plasmid pJL1-sfGFP (Li et al., 2018; Pedelacq et al., 2006) (**Sequence S1**) via an inverse PCR using DNA oligos G302-f and G302-r. As a result, the final LET had the same length and nucleotide sequence as the original plasmid. LET2 was prepared by amplifying the full length of the pJL1-sfGFP plasmid via inverse PCR using DNA oligos G350-f and G350-r. ScCro protein-binding sites with the ORC sequence (<u>TATCACCG</u>CGG<u>GGTGATA</u>) were introduced at the free end of the PCR products by G350-f and G350-r. LET3 was prepared by amplifying the sfGFP expression cassette from the pJL1-sfGFP plasmid via a PCR using DNA oligos G369-f and G369-r, which annealed to the T7 promoter and T7 terminator, thus the PCR product does not carry long buffering sequences at either upstream or downstream regions of the expression cassette. In this way, the LET3 sequence only presents scCro binding sites at the free ends. LET4 was prepared by amplifying the sfGFP expression cassette from the plasmid pJL1-sfGFP via a PCR using the DNA oligos G370-f and G370-r. LET5 was prepared by amplifying the sfGFP expression cassette via a PCR using the DNA oligos G243-f and G243-r that contain three phosphorothioate bonds modifications at the 5’ ends to prevent the hydrolysis of phosphodiester bonds caused by other exonucleases (Putney et al., 1981; Yang et al., 2007) outside of RecBCD. The PCR products mentioned above were resolved on 1.2 % agarose gel (Invitrogen) and purified by the QIAquick PCR Purification Kit (Qiagen). All DNA samples were quantified by the Nanodrop 2000c spectrophotometer (Thermo Scientific).

### LET methylation

The LET2 was methylated with dam or CpG methyltransferases following the manufacturer’s instructions. The reactions were then purified by the QIAquick PCR Purification Kit (Qiagen) and quantified using a Nanodrop 2000c spectrophotometer.

### scCro protein expression and purification

The scCro expression plasmid pET22-scCro previously described (Kojima et al., 2016) was transformed into the *E. coli* strain BL21(DE3). A 100 mL of 2 × YT medium containing 50 µg/mL carbenicillin was inoculated with 1 mL of overnight pre-culture. Cells were grown at 37 °C to OD_600_ ∼ 0.5, then the expression of scCro was induced with 1 mM IPTG for 5 hours at 37 °C. Next, cells were harvested by centrifugation at 6,000 × g for 10 min at 4 °C, and stored at −25 °C. The frozen pellet (from 50 mL of culture) was thawed and suspended in 2 mL (6 mL per gram of wet pellet) of binding buffer (50 mM sodium phosphate, 300 mM KCl, 10 mM imidazole and 1.4 mM 2-mercaptoethanol, pH 8) containing 1 mM phenylmethylsulfonyl fluoride and 1 mg/mL of lysozyme. A 1 mL cell suspension was incubated at 4 °C for 30 min and ultrasonicated on ice (30 s × 2, approx. 300 J) by Q125 Sonicator (Qsonica) with a 3.175 mm diameter probe at a frequency of 20 kHz and 50 % of amplitude. The lysate was recovered by centrifugation at 12,000 × g for 20 min at 4 °C. Following lysate recovery, 900 μL of the supernatant was passed through an Ni-NTA spin-down column (Qiagen) according to the manufacturer’s instructions. The column was then washed three times with 450 μL of washing buffer (50 mM sodium phosphate, 300 mM KCl, 50 mM imidazole and 1.4 mM 2-mercaptoethanol, pH 8). Proteins bound to the resin were eluted in 4 fractions with 450 μL of elution buffer (50 mM sodium phosphate, 300 mM KCl, 500 mM imidazole and 1.4 mM 2-mercaptoethanol, pH 8). Elution fractions were assessed by SDS-PAGE, and the E3 and E4 fractions were combined and dialyzed twice against 500 mL of S30 buffer 2 (10 mM Tris-OAc, 14 mM Mg(OAc)_2_, 60 mM KOAc, 1 mM DTT, pH 8.2) and once against 500 mL of S30 buffer 2 containing 50 % glycerol. Finally, the concentration of purified scCro was determined using the Quick Start™ Bradford Protein Assay Kit (Bio-Rad) and the sample was stored at -25 °C (**Figure S3**).

### scCro protection prevents RecBCD degradation of LETs

The RecBCD degradation test was performed following the instructions provided by New England Biolabs (NEB) with a minor modification detailed as follows. LET2 and scCro were pre-mixed in a 1.5 µL mixture at room temperature for 60 min to allow binding of scCro protein to linear DNA templates and added to a 10 µL of RecBCD degradation reaction containing 1x NEB buffer 4, 1 mM of ATP, and 0.2 U/µL of RecBCD. For the time course study, samples were harvested at 15 min intervals, and the reaction was immediately terminated by diluting with an equal volume of 30 mM EDTA solution followed by a phenol-chloroform extraction. The mixture was centrifuged for 1 min at maximum speed, and 10 µL of the supernatant was recovered and resolved using a 1 % agarose gel.

### sfGFP expression in CFPS from LETs

Expression and quantification of sfGFP in *E. coli*-based CFPS was performed according to a previously published report with minor modification (Hong et al., 2014; Kwon and Jewett, 2015). Briefly, the C321.ΔA.759 (*endA*^−^ *gor*^−^ *rne*^−^ *mazF*^−^)(Martin et al., 2018) *E. coli* strain was used for preparing the S12 extract. The purified scCro and the plasmid or LETs mixture was incubated at room temperature for 1 hr before adding into the CFPS reaction. The final concentrations of scCro and expression template were 2 μM and 8 nM, respectively, unless stated otherwise. The GamS-based inhibition of RecBCD was performed according to a previously published report (Sun et al., 2014).

The *V. natriegens* S30 extract preparation and CFPS reaction was performed according to previous publication without any modification (Des Soye et al., 2018). For protection, LET was mixed with scCro protein in a 3-μL volume for binding at room temperature for 1 hour. In parallel, non-protected LET was prepared in the same way except no scCro supplied. Thereafter, these pretreated LETs were directly added to a final 15 μL of *V. natriegens* CFPS for a 16-hour incubation at 30 °C. Plasmid pJL1-sfGFP was used as a circular template control. In *V. natriegens* CFPS, LET was supplied at four different final concentrations: 8 nM, 16 nM, 32 nM, and 64 nM while the final concentration of scCro protein was fixed at 2 μM for all samples.

## Acknowledgments

We thank graduate student Weston Kightlinger in the Jewett Lab for preparation of the GamS protein. We also thank Ashty Karim for his input on the manuscript. This work was supported in part by a Grant-in-Aid (no. 19H02523) from the Ministry of Education, Culture, Sports, Science, and Technology of Japan (MEXT) to H.N.; M.C.J. acknowledges support from the DARPA 1000 Molecules Program HR0011-15-C-0084, Army Research Office Grants W911NF-16-1-0372, W911NF-19-1-0298, and W911NF-18-1-0200; the National Institutes of Health Grant 1U19AI142780-01, the David and Lucile Packard Foundation, and the Camille Dreyfus Teacher-Scholar Program.

## Author Contributions

B.Z., R.G., T.K., M.C.J. and H.N. designed the experiments. B.Z. and R.G. conducted the experiments. M.D.C. prepared *V. natriegens* cell-free extract. B.Z., R.G., M.C.J. and H.N. wrote the paper. All authors discussed the results and edited the manuscript.

## Abbreviations

LET: linear expression template;
CFPS: cell-free protein synthesis;
sfGFP: superfolder green fluorescent protein;
scCro: single-chain derivative of the lambda Cro repressor

## References

Albright, R. A., & Matthews, B. W. (1998). Crystal structure of lambda-Cro bound to a consensus operator at 3.0 A resolution. Journal of Molecular Biology, 280, 137–151. doi: 10.1006/jmbi.1998.1848

Carlson, E. D., Gan, R., Hodgman, C. E., & Jewett, M. C. (2012). Cell-free protein synthesis: applications come of age. Biotechnology Advances, 30, 1185–1194. doi: 10.1016/j.biotechadv.2011.09.016

Des Soye, B. J., Davidson, S. R., Weinstock, M. T., Gibson, D. G., & Jewett, M. C. (2018). Establishing a high-yielding cell-free protein synthesis platform derived from *Vibrio natriegens*. ACS Synthetic Biology, 7, 2245–2255. doi: 10.1021/acssynbio.8b00252

Des Soye, B. J., Gerbasi, V. R., Thomas, P. M., Kelleher, N. L., & Jewett, M. C. (2019). A highly productive, one-pot cell-free protein synthesis platform based on genomically recoded *Escherichia coli*. Cell Chemical Biology, 26, 1743–1754 e1749. doi: 10.1016/j.chembiol.2019.10.008

Failmezger, J., Scholz, S., Blombach, B., & Siemann-Herzberg, M. (2018). Cell-free protein synthesis from fast-growing Vibrio natriegens. Frontiers in Microbiology, 9, 1146. doi: 10.3389/fmicb.2018.01146

Harris, D. C., & Jewett, M. C. (2012). Cell-free biology: Exploiting the interface between synthetic biology and synthetic chemistry. Current Opinion in Biotechnology, 23, 672–678. doi: 10.1016/j.copbio.2012.02.002

Hoffmann, T., Nemetz, C., Schweizer, R., Mutter, W., & Watzele, M. (2002). High-level cell-free protein expression from PCR-generated DNA templates. In: Spirin, A. S., editor. Cell-free Translation Systems. Berlin, Heidelberg: Springer. pp. p203–210.

Hong, S. H., Ntai, I., Haimovich, A. D., Kelleher, N. L., Isaacs, F. J., & Jewett, M. C. (2014). Cell-free protein synthesis from a release factor 1 deficient *Escherichia coli* activates efficient and multiple site-specific nonstandard amino acid incorporation. ACS Synthetic Biology, 3, 398–409. doi: 10.1021/sb400140t

Jana, R., Hazbun, T. R., Fields, J. D., & Mossing, M. C. (1998). Single-chain lambda Cro repressors confirm high intrinsic dimer-DNA affinity. Biochemistry, 37, 6446–6455. doi: 10.1021/bi980152v

Karim, A. S., & Jewett, M. C. (2016). A cell-free framework for rapid biosynthetic pathway prototyping and enzyme discovery. Metabolic Engineering, 36, 116–126. doi: 10.1016/j.ymben.2016.03.002

Kelwick, R., Webb, A. J., MacDonald, J. T., & Freemont, P. S. (2016). Development of a *Bacillus subtilis* cell-free transcription-translation system for prototyping regulatory elements. Metabolic Engineering, 38, 370–381. doi: 10.1016/j.ymben.2016.09.008

Kightlinger, W., Duncker, K. E., Ramesh, A., Thames, A. H., Natarajan, A., Stark, J. C., … Jewett, M. C. (2019). A cell-free biosynthesis platform for modular construction of protein glycosylation pathways. Nature Communications, 10, 5404. doi: 10.1038/s41467-019-12024-9

Kim, J. G., Takeda, Y., Matthews, B. W., & Anderson, W. F. (1987). Kinetic studies on Cro repressor-operator DNA interaction. Journal of Molecular Biology, 196, 149–158. doi: 10.1016/0022-2836(87)90517-1

Kojima, T., Mizoguchi, T., Ota, E., Hata, J., Homma, K., Zhu, B., … Nakano, H. (2016). Immobilization of proteins onto microbeads using a DNA binding tag for enzymatic assays. Journal of Bioscience and Bioengineering, 121, 147–153. doi: 10.1016/j.jbiosc.2015.06.003

Kojima, T., Hata, J., Oka, H., Hayashi, K., Hitomi, K., & Nakano, H. (2018). Spatial arrangement of proteins using scCro-tag: application for an *in situ* enzymatic microbead assay. Bioscience, Biotechnology, and Biochemistry, 82, 1911–1921. doi: 10.1080/09168451.2018.1501265

Kwon, Y. C., & Jewett, M. C. (2015). High-throughput preparation methods of crude extract for robust cell-free protein synthesis. Scientific Reports, 5, 8663. doi: 10.1038/srep08663

Li, J., Wang, H., Kwon, Y. C., & Jewett, M. C. (2017). Establishing a high yielding *streptomyces*-based cell-free protein synthesis system. Biotechnology and Bioengineering, 114, 1343–1353. doi: 10.1002/bit.26253

Li, J., Wang, H., & Jewett, M. C. (2018). Expanding the palette of *Streptomyces*-based cell-free protein synthesis systems with enhanced yields. Biochemical Engineering Journal, 130, 29–33. doi: 10.1016/j.bej.2017.11.013

Lin, L., Kightlinger, W., Prabhu, S. K., Hockenberry, A. J., Li, C., Wang, L. X.,… Mrksich, M. (2020). Sequential glycosylation of proteins with substrate-specific *N*-glycosyltransferases. ACS Central Science, 6, 144–154. doi: 10.1021/acscentsci.9b00021

Marshall, R., Maxwell, C. S., Collins, S. P., Beisel, C. L., & Noireaux, V. (2017). Short DNA containing chi sites enhances DNA stability and gene expression in *E. coli* cell-free transcription-translation systems. Biotechnology and Bioengineering, 114, 2137–2141. doi: 10.1002/bit.26333

Martin, R. W., Majewska, N. I., Chen, C. X., Albanetti, T. E., Jimenez, R. B. C., Schmelzer, A. E., … Roy, V. (2017). Development of a CHO-based cell-free platform for synthesis of active monoclonal antibodies. ACS Synthetic Biology, 6, 1370–1379. doi: 10.1021/acssynbio.7b00001

Martin, R. W., Des Soye, B. J., Kwon, Y. C., Kay, J., Davis, R. G., Thomas, P. M., … Jewett, M. C. (2018). Cell-free protein synthesis from genomically recoded bacteria enables multisite incorporation of noncanonical amino acids. Nature Communications, 9, 1203. doi: 10.1038/s41467-018-03469-5

Michel-Reydellet, N., Woodrow, K., & Swartz, J. (2005). Increasing PCR fragment stability and protein yields in a cell-free system with genetically modified *Escherichia coli* extracts. Journal of Molecular Microbiology and Biotechnology, 9, 26–34. doi: 10.1159/000088143

Nilsson, M. T., & Widersten, M. (2004). Repertoire selection of variant single-chain Cro: toward directed DNA-binding specificity of helix-turn-helix proteins. Biochemistry, 43, 12038–12047. doi: 10.1021/bi049122k

Pedelacq, J. D., Cabantous, S., Tran, T., Terwilliger, T. C., & Waldo, G. S. (2006). Engineering and characterization of a superfolder green fluorescent protein. Nature Biotechnology, 24, 79–88. doi: 10.1038/nbt1172

Putney, S. D., Benkovic, S. J., & Schimmel, P. R. (1981). A DNA fragment with an alpha-phosphorothioate nucleotide at one end is asymmetrically blocked from digestion by exonuclease III and can be replicated in vivo. Proceedings of the National Academy of Sciences of the United States of America, 78, 7350–7354. doi: 10.1073/pnas.78.12.7350

Salzberg, S. L. (2019). Next-generation genome annotation: we still struggle to get it right. Genome Biology, 20, 92. doi: 10.1186/s13059-019-1715-2

Shrestha, P., Smith, M. T., & Bundy, B. C. (2014). Cell-free unnatural amino acid incorporation with alternative energy systems and linear expression templates. New Biotechnology, 31, 28–34. doi: 10.1016/j.nbt.2013.09.002

Silverman, A. D., Karim, A. S., & Jewett, M. C. (2020). Cell-free gene expression: an expanded repertoire of applications. Nature Reviews Genetics, 21, 151–170. doi: 10.1038/s41576-019-0186-3

Stark, J. C., Huang, A., Hsu, K. J., Dubner, R. S., Forbrook, J., Marshalla, S., … Jewett, M. C. (2019). BioBits Health: Classroom activities exploring engineering, biology, and human health with fluorescent readouts. ACS Synthetic Biology, 8, 1001–1009. doi: 10.1021/acssynbio.8b00381

Sun, Z. Z., Yeung, E., Hayes, C. A., Noireaux, V., & Murray, R. M. (2014). Linear DNA for rapid prototyping of synthetic biological circuits in an *Escherichia coli* based TX-TL cell-free system. ACS Synthetic Biology, 3, 387–397. doi: 10.1021/sb400131a

Swartz, J. R., Jewett, M. C., & Woodrow, K. A. (2004). Cell-free protein synthesis with prokaryotic combined transcription-translation. In: P., B., and A., L., editors. Recombinant Gene Expression. Methods in Molecular Biology. Totowa, New Jersey: Humana Press. pp. p169–182.

Thavarajah, W., Silverman, A. D., Verosloff, M. S., Kelley-Loughnane, N., Jewett, M. C., & Lucks, J. B. (2020). Point-of-use detection of environmental fluoride via a cell-free riboswitch-based biosensor. ACS Synthetic Biology, 9, 10–18. doi: 10.1021/acssynbio.9b00347

Ueno, S., Yoshida, S., Mondal, A., Nishina, K., Koyama, M., Sakata, I., … Sakai, T. (2012). *In vitro* selection of a peptide antagonist of growth hormone secretagogue receptor using cDNA display. Proceedings of the National Academy of Sciences of the United States of America, 109, 11121–11126. doi: 10.1073/pnas.1203561109

Wang, H., Li, J., & Jewett, M. C. (2018). Development of a *Pseudomonas putida* cell-free protein synthesis platform for rapid screening of gene regulatory elements. Synthetic Biology, 3, ysy003. doi: 10.1093/synbio/ysy003

Wiegand, D. J., Lee, H. H., Ostrov, N., & Church, G. M. (2018). Establishing a cell-free *Vibrio natriegens* expression system. ACS Synthetic Biology, 7, 2475–2479. doi: 10.1021/acssynbio.8b00222

Yang, Z. Y., Sismour, A. M., & Benner, S. A. (2007). Nucleoside alpha-thiotriphosphates, polymerases and the exonuclease III analysis of oligonucleotides containing phosphorothioate linkages. Nucleic Acids Research, 35, 3118–3127. doi: 10.1093/nar/gkm168

Zhu, B., Mizoguchi, T., Kojima, T., & Nakano, H. (2015). Ultra-high-throughput screening of an *in vitro*-synthesized horseradish peroxidase displayed on microbeads using cell sorter. PLOS ONE, 10. doi: 10.1371/journal.pone.0127479

