## Supplementary data and figures for "Increasing cell-free gene expression yields from linear templates in *Escherichia coli* and *Vibrio natriegens* extracts by using DNA-binding proteins"

### *Supplementary Material*

Supplemental Figure S1, describing the effects of GamS and scCro on improving yield of the *E. coli* CFPS platform; Supplemental Figure S2, shows the optimization of scCro concentrations in *E. coli* CFPS reaction; Supplemental Figure S3, shows the SDS-PAGE for the purification of the scCro protein; Supplemental Table S1 listing the sequences of the oligonucleotides used in this study; Supplemental Sequence S1 listing the sequences of the pJL1-sfGFP plasmid.

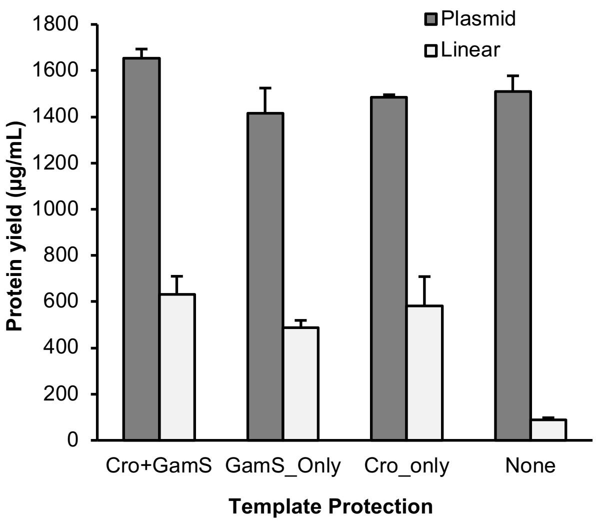


**Figure S1. Comparison of scCro-based protection and GamS-based inhibition on increasing the sfGFP yield.** Plasmid indicates the samples using plasmid DNA as the expression template. Linear indicates the samples using LET2 as the expression template. The mean and standard deviations are shown (N=4).

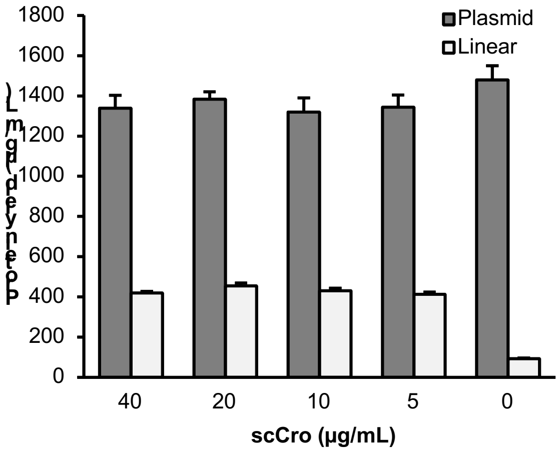


**Figure S2. Effect of concentration of scCro on the sfGFP yield.** Plasmid indicates the samples using plasmid DNA as the expression template. Linear indicates the samples using LET2 as the expression template. The mean and standard deviations are shown (N=4). Molecular weight of scCro is 17.7 kDa. The scCro molar concentration ranges from 5 to 40 μg/mL (0.28 – 2.2 μM) in this test.

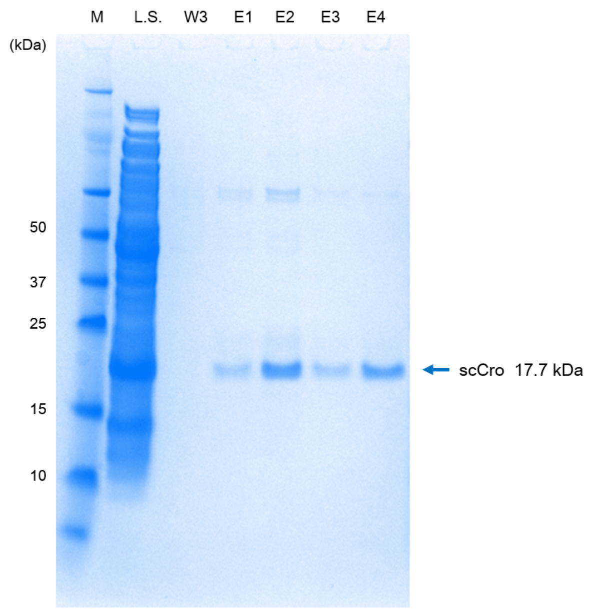


**Figure S3. SDS-PAGE of purified scCro using $Ni^{2+}$-affinity chromatography.** M: protein marker; L.S.: lysate supernatant; W3: final wash fraction; E1-E4: elution fractions 1through 4. The blue arrow corresponds to the molecular weight of scCro (17.7 kDa).

**Table S1: List of DNA oligonucleotides used in this study.**
Asterisk after a base indicates a phosphorothioate bond modification.

| Name | Sequence |
|---|---|
| G243-f | G*A*A*TTCAGATCTCGATCCCGCGAAATTAATACGACTCACTATAGG |
| G243-r | C*C*C*GTTTAGAGGCCCCAAGGGGTTATGCTAGTTATTGCTCAGCGG |
| G302-f | GATAGATTGTCGCACCTGATT |
| G302-r | GATTGTATGGGAAGCCCGATG |
| G350-f | GATCCTATCACCGCGGGTGATAGTACGGATAGATTGTCGCACCTGATT |
| G350-r | GATCCTATCACCGCGGGTGATAGTACGGATTGTATGGGAAGCCCGATG |
| G369-f | GATCCTATCACCGCGGGTGATAGTACGGATCCCGCGAAATTAATACGACTCACTATAGG |
| G369-r | GATCCTATCACCGCGGGTGATAGTACGCCCCAAGGGGTTATGCTAGTTATTGCTCAGCGG |
| G370-f | GTACGGATCCCGCGAAATTAATACGACTCACTATAGG |
| G370-r | GTACGCCCCAAGGGGTTATGCTAGTTATTGCTCAGCGG |

**Sequence S1: Full DNA sequence of pJL1-sfGFP plasmid.**

>pJL1-sfGFP, KanR, sfGFP gene, pBR322 ori, 2486 bp

```
agatcaaaggatcttcttgagatcctttttttctgcgcgtaatctgctgcttgcaaacaaaaaa
accaccgctaccagcggtggtttgtttgccggatcaagagctaccaactctttttccgaaggta
actggcttcagcagagcgcagataccaaatactgttcttctagtgtagccgtagttaggccacc
acttcaagaactctgtagcaccgcctacatacctcgctctgctaatcctgttaccagtggctgc
tgccagtggcgataagtcgtgtcttaccgggttggactcaagacgatagttaccggataaggcg
cagcggtcgggctgaacggggggttcgtgcacacagcccagcttggagcgaacgacctacaccg
aactgagatacctacagcgtgagctatgagaaagcgccacgcttcccgaagggagaaaggcgga
caggtatccggtaagcggcagggtcggaacaggagagcgcacgagggagcttccagggggaaac
gcctggtatctttatagtcctgtcgggtttcgccacctctgacttgagcgtcgatttttgtgat
gctcgtcaggggggcggagcctatggaaaaacgccagcaacgcgatcccgcgaaattaatacga
ctcactatagggagaccacaacggtttccctctagaaataattttgtttaactttaagaaggag
atatacatatgagcaaaggtgaagaactgtttaccggcgttgtgccgattctggtggaactgga
tggcgatgtgaacggtcacaaattcagcgtgcgtggtgaaggtgaaggcgatgccacgattggc
aaactgacgctgaaatttatctgcaccaccggcaaactgccggtgccgtggccgacgctggtga
ccaccctgacctatggcgttcagtgttttagtcgctatccggatcacatgaaacgtcacgattt
ctttaaatctgcaatgccggaaggctatgtgcaggaacgtacgattagctttaaagatgatggc
aaatataaaacgcgcgccgttgtgaaatttgaaggcgataccctggtgaaccgcattgaactga
aaggcacggattttaaagaagatggcaatatcctgggccataaactggaatacaactttaatag
ccataatgtttatattacggcggataaacagaaaaatggcatcaaagcgaattttaccgttcgc
cataacgttgaagatggcagtgtgcagctggcagatcattatcagcagaataccccgattggtg
atggtccggtgctgctgccggataatcattatctgagcacgcagaccgttctgtctaaagatcc
gaacgaaaaaggcacgcgggaccacatggttctgcacgaatatgtgaatgcggcaggtattacg
tggagccatccgcagttcgaaaaataagtcgaccggctgctaacaaagcccgaaaggaagctga
gttggctgctgccaccgctgagcaataactagcataaccccttggggcctctaaacgggtcttg
aggggttttttgctgaaagccaattctgattagaaaaactcatcgagcatcaaatgaaactgca
atttattcatatcaggattatcaataccatatttttgaaaaagccgtttctgtaatgaaggaga
aaactcaccgaggcagttccataggatggcaagatcctggtatcggtctgcgattccgactcgt
ccaacatcaatacaacctattaatttcccctcgtcaaaaataaggttatcaagtgagaaatcac
catgagtgacgactgaatccggtgagaatggcaaaagcttatgcatttctttccagacttgttc
aacaggccagccattacgctcgtcatcaaaatcactcgcatcaaccaaaccgttattcattcgt
gattgcgcctgagcgagacgaaatacgcgatcgctgttaaaaggacaattacaaacaggaatcg
aatgcaaccggcgcaggaacactgccagcgcatcaacaatattttcacctgaatcaggatattc
ttctaatacctggaatgctgttttcccggggatcgcagtggtgagtaaccatgcatcatcagga
gtacggataaaatgcttgatggtcggaagaggcataaattccgtcagccagtttagtctgacca
tctcatctgtaacatcattggcaacgctacctttgccatgtttcagaaacaactctggcgcatc
gggcttcccatacaatcgatagattgtcgcacctgattgcccgacattatcgcgagcccattta
tacccatataaatcagcatccatgttggaatttaatcgcggcttcgagcaagacgtttcccgtt
gaatatggctcataacaccccttgtattactgtttatgtaagcagacagttttattgttcatga
tgatatatttttatcttgtgcaatgtaacatcagagattttgagacacaacgtg
```
